# Effect of Acute Alcohol Consumption in a Novel Rodent Model of Decision Making

**DOI:** 10.1101/2024.09.24.614857

**Authors:** Atanu Giri, Serina A. Batson, Andrea Y. Macias, Cory N. Heaton, Neftali F. Reyes, Alexis A. Salcido, Luis D. Davila, Lara I. Rakocevic, Dirk W. Beck, Raquel J. Ibañez Alcalá, Safa B. Hossain, Paulina Vara, Sabrina M. Drammis, Kenichiro Negishi, Adrianna E. Rosales, Laura E. O’Dell, Travis M. Moschak, Ki A. Goosens, Alexander Friedman

## Abstract

**Background:** Alcohol use, especially at high consumption levels, can lead to irrational decision-making. In humans, this can lead to harmful outcomes often seen in the context of driving under the influence and or aggressive behavior. To date, the field is lacking comprehensive animal models to examine the impact of alcohol use on decision making in rodents, particularly to examine sex differences in choice behavior. To address this issue, the present study examined the effects of acute alcohol consumption during a behavioral approach-avoidance task that captures momentary changes in decision-making behavior and choice selection in female and male rats.

**Methods:** Our team has developed a novel behavioral protocol involving a concurrent choice to consume four different concentrations of alcohol and sucrose combinations. During the task, female or male rats can approach or avoid drinking solutions in four distinct corners of our test apparatus. The solutions were prepared in inverse concentrations (higher sucrose was paired with lower alcohol and vice versa) so that the rodents pursue minimal alcohol use by consuming the higher sucrose concentrations or higher concentrations of alcohol by drinking the lower sucrose concentrations. The animals also have the option to avoid drinking alcohol by not approaching any of the drinking cups. Behavior and choice were tracked during task performance involving different solution concentrations of alcohol and sucrose.

**Results:** The choice of consuming different concentrations of alcohol or sucrose resulted in sex-dependent differences in an approach-avoid trade-off pattern of behavior that was sensitive to different concentrations of alcohol/sucrose combinations. Notably, males were greatly affected by the introduction of alcohol into the task environment, approaching higher alcohol concentrations significantly more often than the non-alcohol containing options. In contrast, females choice patterns and task performance were largely unchanged during alcohol and non-alcohol containing tasks. Regardless of sex, we identify a novel method for identifying individual subject decision-making abnormalities during and after alcohol consumption.

**Conclusions:** This research reveals a novel approach for examining the effects of acute alcohol exposure during a trade-off task, with decision patterns being more impacted by alcohol use in males as compared to females. We also offer the field a novel approach for identifying individual abnormalities in decision making behavior with the presentation of alcohol. Future research can explore these abnormal patterns in both acute and chronic alcohol conditions to develop methods for identifying subjects at-risk for developing an alcohol use disorder and the deleterious impact of alcohol on rational decision making.

## Introduction

Decision-making is a fundamental process for optimal day-to-day functioning. A recent review of the literature concluded that the current value-based decision-making models at the preclinical level treat decision making as a uniform process (Orsini *et al*., 2019). This can be limiting when trying to assess changes in decision-making following exposure to substances, such as alcohol, which have both aversive and rewarding qualities. Decision-making changes are an important metric for gauging the impact alcohol on an individual or across groups that may be impacted by alcohol use to a greater degree, such as females versus males(Flores-Bonilla and Richardson, 2020). Indeed, alcohol consumption can impair day-to-day decision-making (Bechara, 2005), increase risky decision-making (Fein, Klein and Finn, 2004; Lane *et al*., 2004; George, Rogers and Duka, 2005; Noël *et al*., 2007; Bidwell *et al*., 2013; Kornreich *et al*., 2013; Brevers *et al*., 2014; Wallin-Miller *et al*., 2017; Aguirre *et al*., 2020; Burnette *et al*., 2021), and alter other cognitive functions (Weissenborn and Duka, 2003; Field *et al*., 2010; Dry *et al*., 2012; Van Skike, Goodlett and Matthews, 2019). Alcohol is used widely across the globe (Degenhardt *et al*., 2008; Rehm, 2016); thus, it is critical to explore how decision-making is affected by the presentation of alcohol and how subsequent decisions may be influenced by prior alcohol-related choices.

Alcohol consumption can have different effects depending on factors like tolerance, genetics, and states like stress (Weafer and Fillmore, 2008). These factors could all contribute to individual differences in decision-making causing increased impulsivity in some(Weissenborn and Duka, 2003) while minimally affecting others (Hammersley, Finnigan and Millar, 1994). Furthermore, there are sex differences in how people respond to alcohol consumption (Zachry, Johnson and Calipari, 2019), potentially due to the differences in how alcohol is metabolized (Collins *et al*., 1975). Understanding these individual and sex differences is critical for educating the public to be cognizant of how acute alcohol consumption alters even “simple” decisions. A recent review revealed that women drink less, and are less likely to engage in problem drinking, or develop alcohol-related disorders than men(Erol and Karpyak, 2015). The current evidence suggests that both sex and gender-related factors impact the risk for medical problems and alcohol use disorders in men versus women. One human report revealed sex differences in the impact of alcohol and cannabis use in decision processes, with greater panic behavior in females and greater rates of impulsivity in males(Phillips and Ogeil, 2017).

Rodents are typically averse to consuming alcohol (Anderson, Varlinskaya and Spear, 2010; Pautassi *et al*., 2011), which poses challenges for researchers studying alcohol’s effects on behavior. To address this, some researchers employ methods such as breeding strains of rodents that are genetically predisposed to consuming alcohol (Timme *et al*., 2020; Borruto *et al*., 2021; Sauton *et al*., 2021). Others use forced drinking protocols, where alcohol is mixed with a palatable substance and provided in the home cage (Thiele and Navarro, 2014; Mendoza-Ruiz *et al*., 2018). While these methods are effective for studying chronic alcohol exposure and alcohol dependence, there is a need in the field for models that focus on the choice to consume different concentrations of alcohol.

Various models have been employed to study the impact of alcohol on behavior in rodents. Self-administration tasks are often used to study voluntary alcohol consumption; however, this task only offers a limited number of choices linked to an operant response (Beckwith and Czachowski, 2016). The Conditioned Place Preference (CPP) model provides an opportunity to assess choice behavior for a neutral versus alcohol-paired chamber on a test day after several conditioning sessions. However, this model lacks clear and defined time points when the animal makes decisions, making it challenging to analyze specific moments of choice (Lucke-Wold, 2011). The Runway model is effective in assessing motivated behavior; however, it also lacks the ability to assess discrete choice points, complicating the understanding of underlying neuronal mechanisms during decision-making (Pandy and Khan, 2016). The Iowa Gambling Task (IGT) is used to evaluate risk-taking behavior and complements our task by offering additional perspectives on decision-making under uncertain conditions(Spoelder *et al*., 2015). To expand on the current rodent models, the present study aimed to develop a paradigm that combines etiological validity with a task offering distinct choices, like self-administration but with multiple decision points across different contexts. To achieve this, we utilized our RECORD system to create a more comprehensive and versatile model for studying the effects of acute alcohol consumption on decision-making (Ibáñez Alcalá *et al*., 2024). Our task is also designed to produce psychometric functions of choice, allowing for further measurement and analysis of decision-making behavior.

Using our previously published RECORD system (Ibáñez Alcalá *et al*., 2024), we integrated an alcohol paradigm and implemented a behavioral task in which rats engaged in both low- and high-cost cost-benefit decision-making tasks, with and without alcohol exposure. To summarize, we found that alcohol alters the shape of psychometric functions, representative of changing behavioral patterns (**Fig. 1**), sex differences in these decision-making tasks (**Figs. 2**), and individual variability in response to acute alcohol consumption (**Fig. 3**). We also explore the long-term effects of alcohol on cost-benefit tasks (**Fig. 4**), demonstrating that these effects are not gender-dependent (**Fig. 5**). Additionally, we delve into individual differences in response to long-term alcohol exposure (**Fig. 6**).

**Fig 1:**
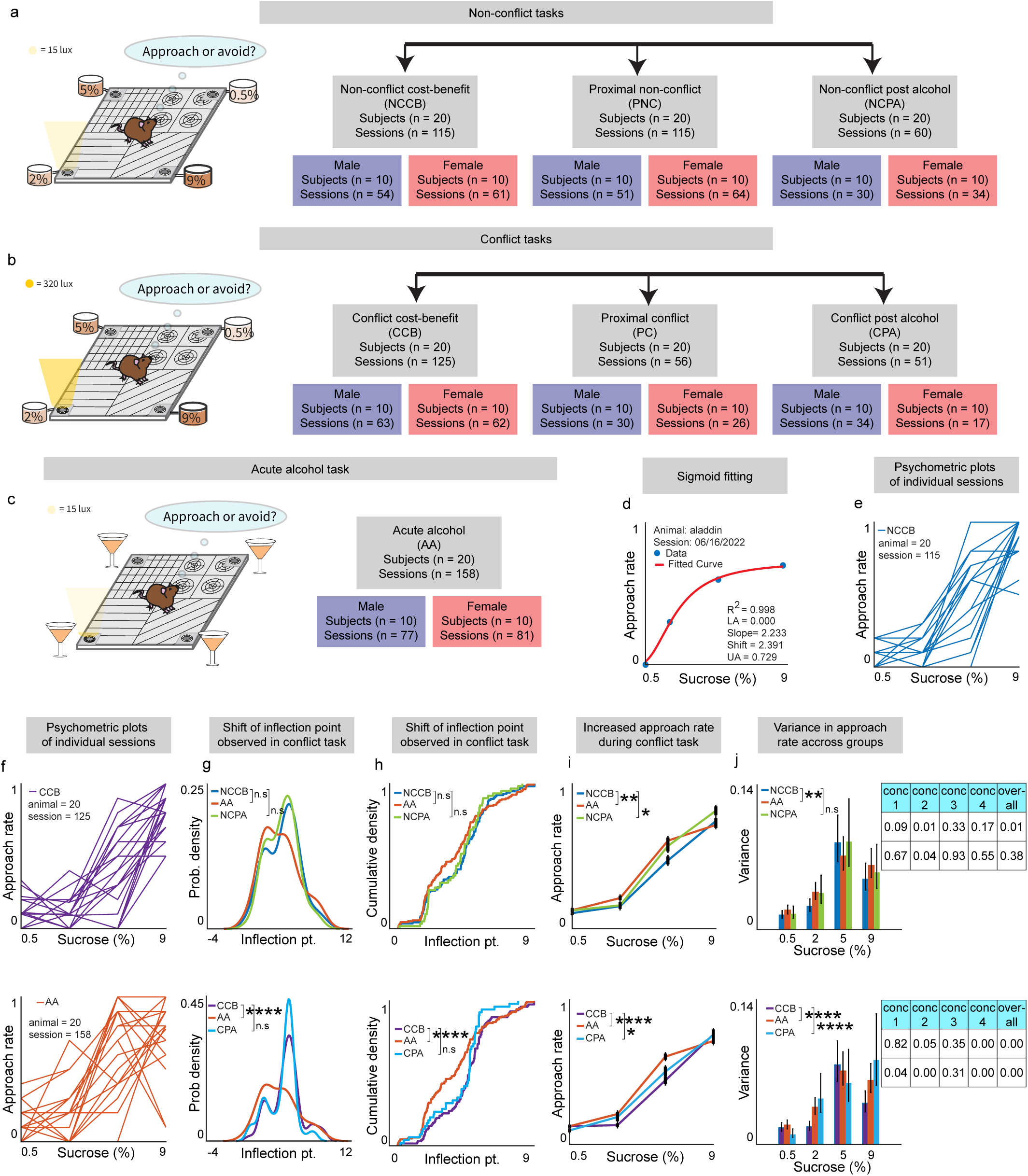
Overview and general analysis of behavioral tasks. (**a**) Non-conflict, (**b**) conflict, and (**c**) acute alcohol cost-benefit tasks utilizing the RECORD setup where one of four sucrose concentrations are dispensed into a bowl surrounded by LED lights. (**d**) Example of psychometric function from a singular non-conflict cost benefit (NCCB) session, fitted with a 4-parameter logistic model (f(x) = d + (a-d)/ (1 + (x/c) ^b)). Individual (n=20) psychometric functions demonstrating approach rate in non-conflict (**e**), conflict (**f, top**), and acute alcohol cost benefit tasks (**f, bottom**). (**g, top**) No significant difference in probability density of inflection points between acute alcohol (AA) compared to NCCB (p = 0.08) or NCPA (p = 0.75). (**g, bottom**) There are significant differences between AA, cost benefit conflict (CCB) tasks (KS test, p < 0.0001), and no significant differences between AA and CPA (p = 0.17). (**h**) Inflection point distribution represented as a cumulative density function. (**i, top**) When comparing NCCB to AA (p = 0.001) and NCPA to NCCB (p = 0.05) approach rates are significantly different. (**i, bottom**) Similarly, comparing CCB to AA (p < 0.0001) and CCB to CPA (p = 0.04) approach rates are also significantly different. (**j**) Variance in approach rate, with error bars depicting upper and lower confidence intervals with 95% significance, show significance between NCCB and AA tasks (F-test, p = 0.0096), and CCB and AA tasks (F-test, p < 0.0001).

**Fig 2:**
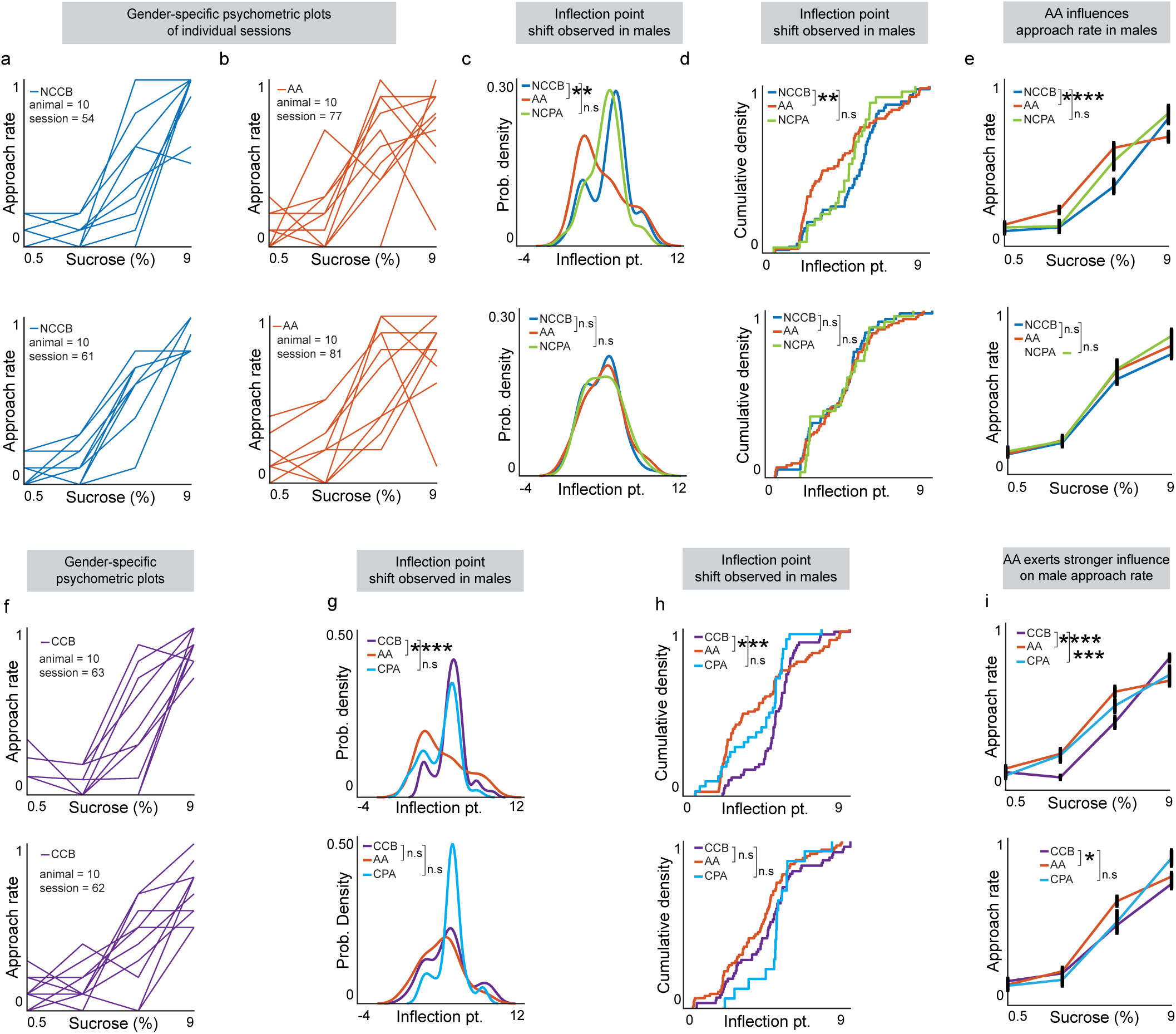
Sex differences during acute alcohol task performance. (**a**) Approach rates for individual sessions for all rats (n=10) in NCCB and (**b**) AA tasks, separated by sex. (**c**) Compared to NCCB, AA inflection point shifted significantly left in males (top, KS test, p = 0.004), but not females (bottom, KS test, p = 0.96). (**d**) There were no significant differences for either sex comparing AA to NCPA (male; p = 0.17, female: p = 0.64). Cumulative density function plots of the distributions in **(c).** (**e**) Approach rate is significantly different between NCCB and AA for males (MANOVA, p< 0.0001, top) but not females (MANOVA, p = 0.47, bottom). (**f**) Approach rates of individual sessions for rats split by sex (n = 10 males, n = 10 females) in CCB. (**g**) Compared to CCB, AA inflection point is significantly shifted left in males (KS test, p < 0.0001) but not in females (KS test, p = 0.211) and again there was so significant difference when comparing CCB to CPA in either sex. (**h**) Inflection point distribution represented as a cumulative density function. (**i**) In CCB vs AA tasks, approach rate is significantly different in males (MANOVA, p < 0.0001) and females (p = 0.0134) in CCB vs AA tasks with males also having significantly approach rates between CCB and CPA (p = 0.0007) unlike females (p = 0.1).

**Fig 3:**
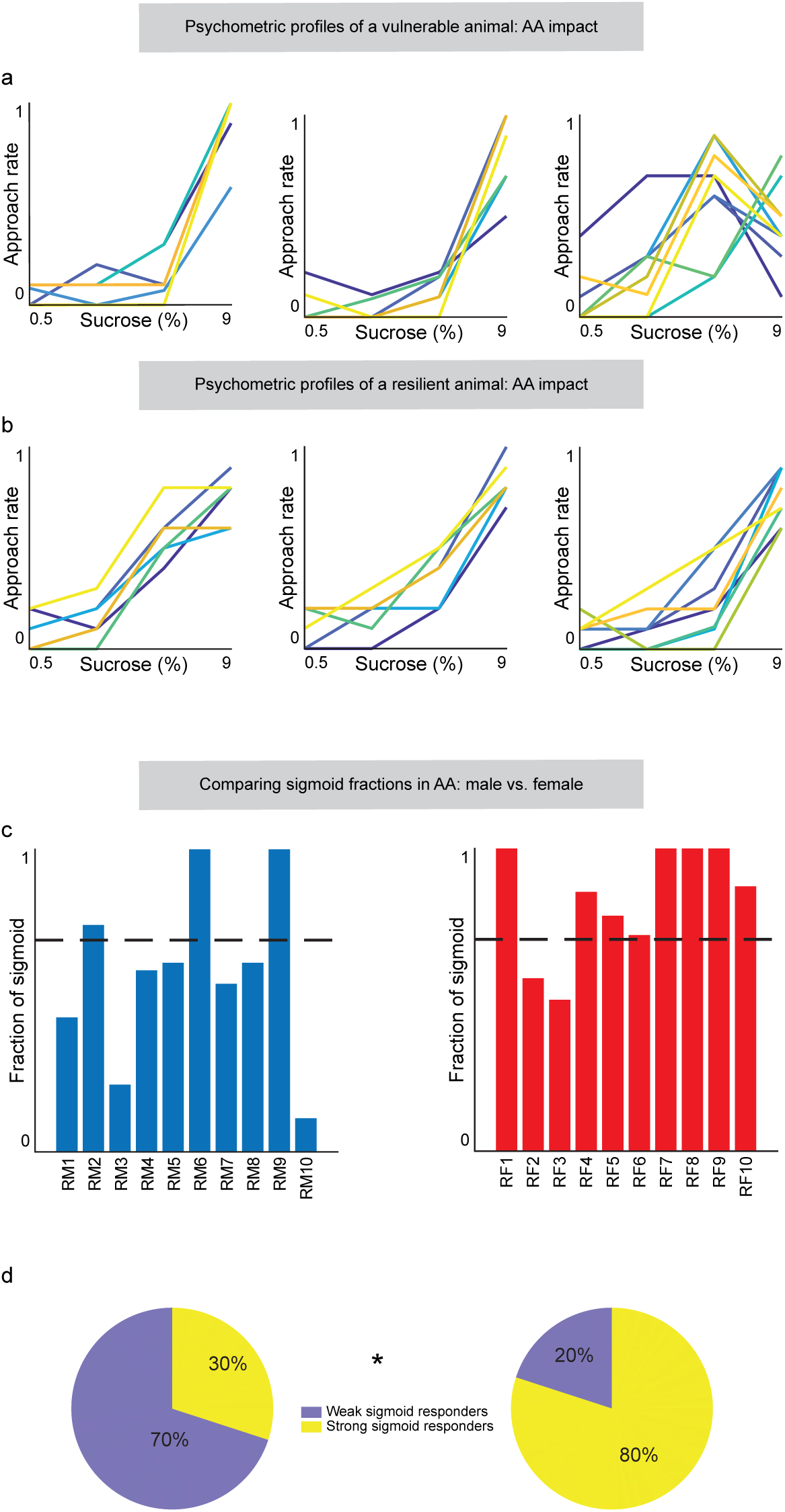
Identifying abnormally performing subjects using their psychometric functions. (**a**) Psychometric profiles depict abnormal task performance of a singular animal during NCCB, CCB, and AA tasks. (**b**) Non-abnormal singular animal psychometric profiles in NCCB, CCB, and AA tasks. (**c**) Two plots showing the fraction of sigmoid across all sessions of AA for individual males (n=10) and females (n=10), where the dashed line at y=0.7 indicates the threshold for fitting a sigmoid psychometric curve. (**d**) Pie charts depicting males (left) and females (right) with sigmoidal (blue) and non-sigmoidal (yellow) psychometric profiles. Males have significantly more non-sigmoidal sessions than females (chi-square test, p = 0.025).

**Fig 4:**
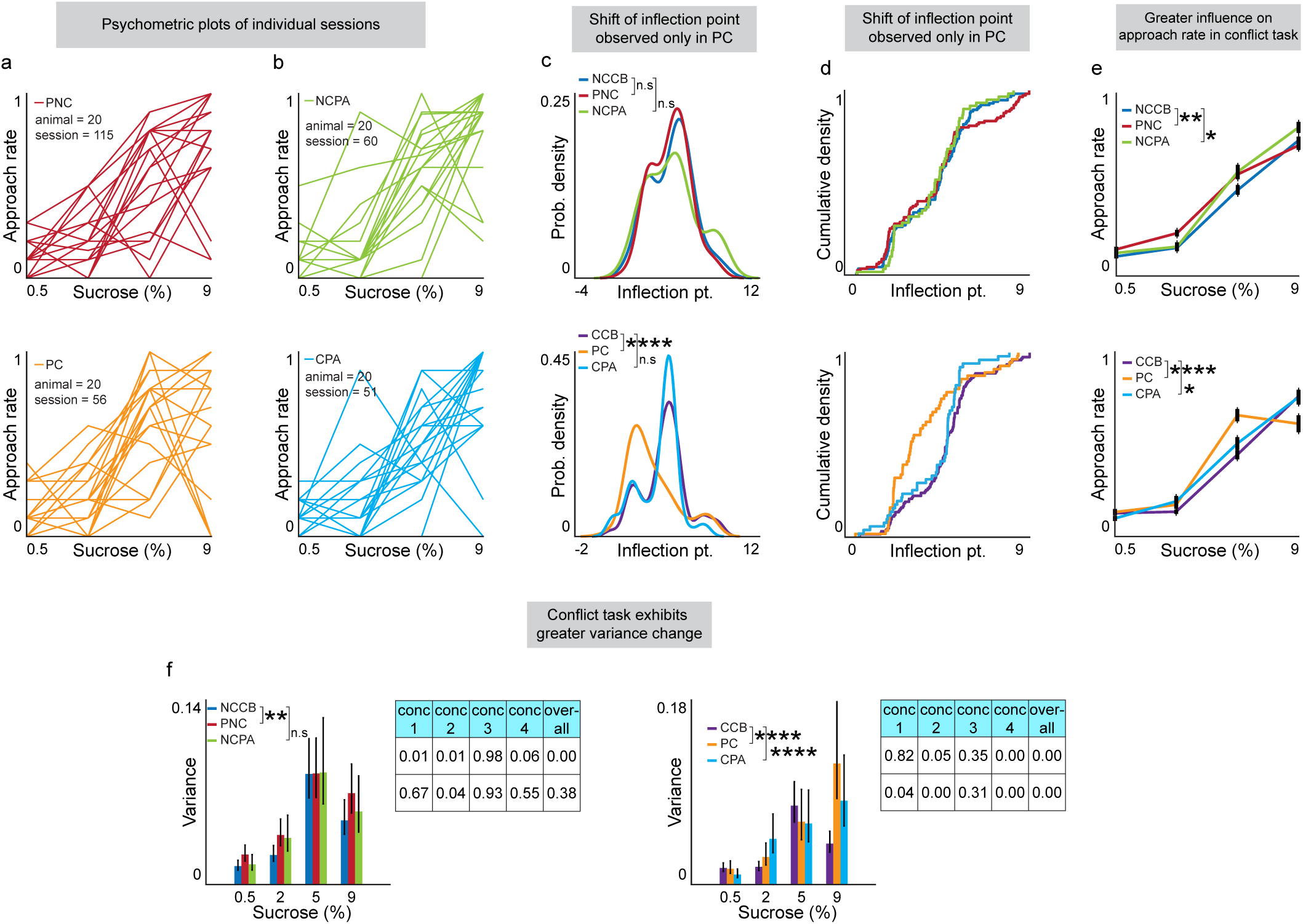
Alcohol effects proximal cost benefit tasks. (**a**) Psychometric plots of approach rates for individual sessions of rats (n=20) in proximal non-conflict (PNC) and proximal conflict (PC) tasks. (**b**) Psychometric plots for individual sessions of all rats (n=20) in non-conflict post-alcohol (NCPA) and conflict post-alcohol (CPA) tasks. (**c**) A significant shift of inflection points in PC compared to CCB (KS test, p=0.0000). (**d**) Inflection point distribution represented as a cumulative density function. (**e**) Approach rates were significantly different across all task types (NCCB vs. PNC: p = 0.002, NCCB vs NCPA; p = 0.05, CCB vs PC; p < 0.0001, CCB vs CPA: 0.04). (**f**) Significant difference in variance between NCCB and PNC in both males (F-test, p = 0.015) and females (F-test, p = 0.0153, **f, left**). Significant variance in CCB vs PC (F-test, p < 0.0001) and CPA (F-test, p < 0.0001) task performance (**f, right**).

**Fig 5:**
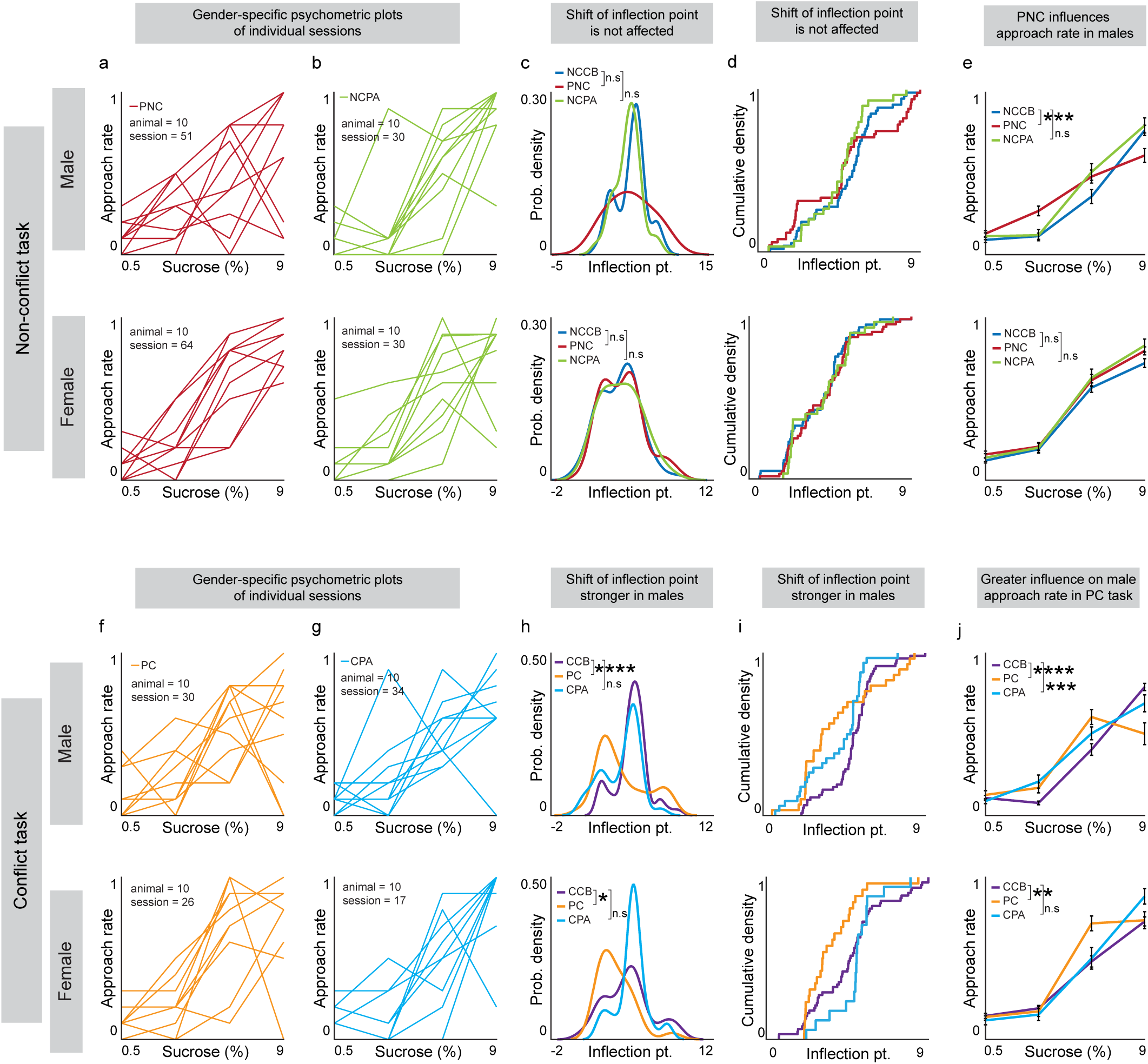
Sex dependent effects of alcohol observed in proximal tasks. Approach rates for individual sessions for all rats (male = 10, female = 10) in (**a**) PNC task and (**b**) NCPA task separated by gender. (**c**) No significant change in inflection point is observed across tasks or genders. (**d**) Inflection point distribution represented as a cumulative density function. (**e**) The difference in approach rates is statistically significant between NCCB and PNC in males (MANOVA, p < 0.0001) but not females (NCCB vs PNC, MANOVA, p = 0.15). Approach rates for individual sessions for all rats (10 males, 10 females) in (**f**) PC and (**g**) CPA, separated by gender. (**h**) PC inflection point is significantly changed compared to CCB in both males (CCB vs. PC: KS test, p = 0.002) and females (CCB vs. PC: KS test, p = 0.02). (**i**) Inflection point distribution represented as a cumulative density function. (**j**) Differences in approach rate is statistically significant between CCB and PC across both males (MANOVA, p = 4.5992e-08) and females (MANOVA, p = 0.0046). Approach rates between CCB and CPA are significant in males (MANOVA, p = 6.5198e-04) but not in females (MANOVA, p = 0.1).

**Fig 6:**
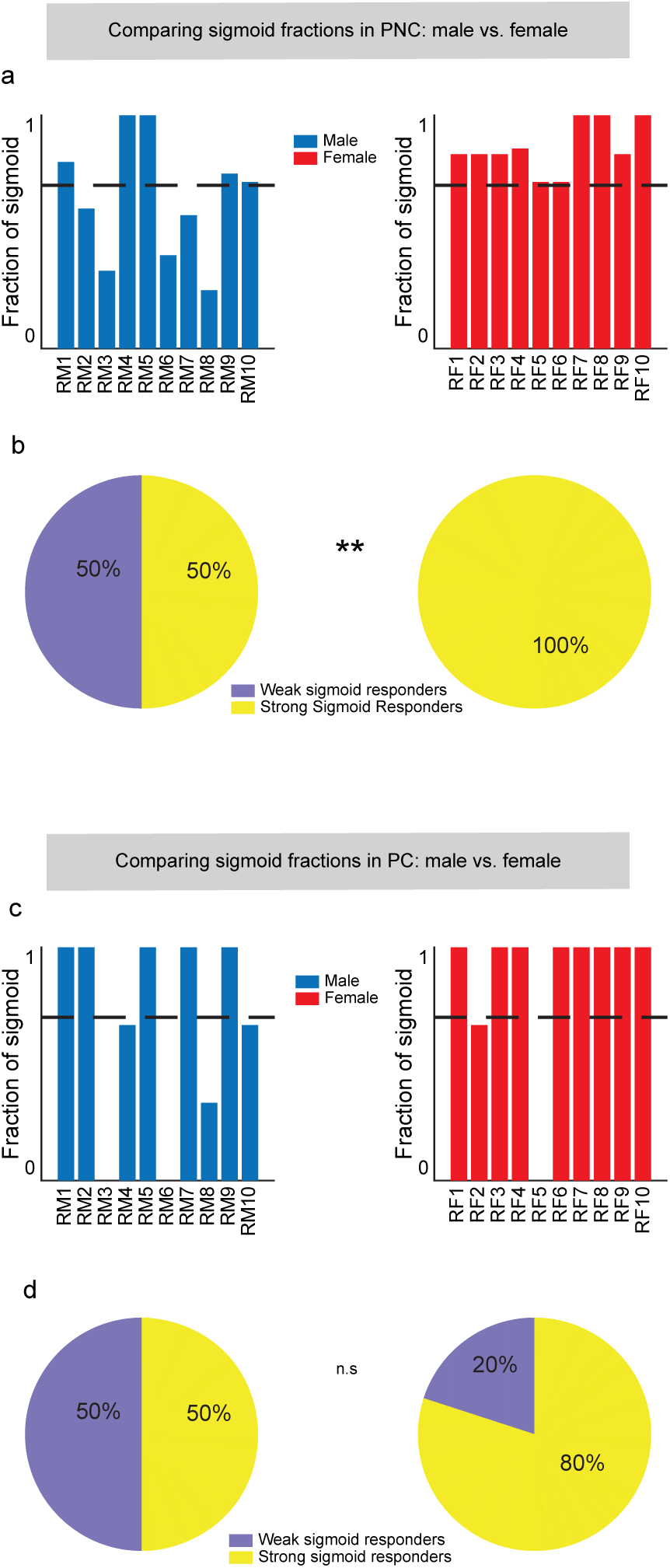
Identification of abnormal performance in proximal tasks. (**a**) Plots showing the fraction of a sigmoid across all sessions from PNC task sessions for individual males (n = 10) and females (n = 10), where the dashed line at y=0.7 represents the threshold for fitting sigmoidal psychometric functions. (**b**) Pie charts showing that there are significantly more abnormal male subjects than females (chi-square test, p = 0.0098) in PNC. (**c**) Plots showing the fraction of sigmoid across all sessions in PC task for individual males (n = 10) and females (n = 10). (**d**) There is no significant difference in the number of abnormal individuals across sex (chi-square test, p = 0.16).

## Materials and Methods

### RECORD framework

We leveraged the RECORD system to introduce alcohol as a component of cost-benefit decision-making within a foraging-like environment (Ibáñez Alcalá *et al*., 2024). On average, training rats to perform RECORD tasks took 9 weeks. The RECORD arena is divided into four quadrants, each distinguished by a different floor pattern (we used diagonal, grid, horizontal, and radial patterns). Rewards were delivered into a “reward zone”, each quadrant had one reward zone and consistently delivered the same sucrose concentration as a reward during tasks (9% diagonal, 5% grid, 2% horizontal, 0.5% radial). LED lights embedded into each reward zone signal where a reward will be delivered with varying brightnesses serving as a cost associated with obtaining the offered reward.

### Non-conflict and conflict tasks

During the non-conflict tasks, the LEDs at each reward zone are set to emit at 15-20 lux. The conflict task, in contrast, sets the LEDs to emit at a maximum of 320-360 lux. During a conflict task, feeders are pseudo-randomly and equally active for low or high-cost brightness for each trial (50% low-brightness: 50% high-brightness). Both tasks are performed in 40-trial sessions, with each trial taking 30 seconds. Signaling the start of a trial, a tone plays for 2.5 seconds. Then one of the reward zones is illuminated and a rat can either approach or avoid the illuminated reward zone. The rat must be in the reward zone for the sucrose reward to be administered otherwise the offer is considered rejected. 4 seconds after reward administration, the LEDs turn off signaling that the trial is over. After 40 trials, rats were returned to their home cage.

### Acute alcohol task

Alcohol was inversely mixed with the preexisting sucrose concentrations and offered with non-conflict task LED brightness (15-20 lux). Solutions consisted of 9 grams of sucrose, 1mL of Everclear brand alcohol (95%), and 99mL of DI water were administered at diagonal part of maze. Solutions containing 5 grams of sucrose, 4mL of Everclear brand alcohol, and 96mL of DI water were administrated at the grid part of the maze. Solutions consisting of 2 grams of sucrose, 10mL of ever clear brand alcohol, and 90mL of DI water were administered at the horizontal part of maze. Lastly, radial reward zones contained 0.5 grams of sucrose, 20mL of Everclear brand alcohol, and 80mL of DI water (Ibáñez Alcalá *et al*., 2024). Experimental sessions, aside from the differences in reward solutions, were performed similarly to the non-conflict task.

### Brief overview of all task types

We used three task types: non-conflict cost benefit task (**Fig 1a**), conflict cost benefit task (**Fig 1b**), and acute alcohol task (**Fig 1c**). The running of these behavioral tasks can be stratified into three major blocks, where numerical order of the block is representative of the order in which tasks were run. Block one, run before alcohol exposure, consists of non-conflict cost benefit (**NCCB**) and conflict cost benefit (**CCB**) tasks repeated across 9 sessions over 4 weeks. Acute alcohol (**AA**) task is performed in experimental block two, where 10 sessions were completed in 5 weeks. Proximal non-conflict (**PNC**) and proximal conflict (**PC**) tasks, defined as running between 1-4 days after AA, also comprise experimental block two. PNC was repeated across 7 sessions over 5 weeks, while 3 PC sessions were completed over 3 weeks. Lastly, block three was performed without alcohol offers in both non-conflict post alcohol (**NCPA**) and conflict post alcohol (**CPA**) tasks for two months after the AA task concluded. NCPA and CPA both ran 3 sessions over 6 weeks. The tasks in each block were run simultaneously, however each animal was limited to one task type per day.

### Spatiotemporal behavioral dynamics

Behavioral features derived from behavioral data, like approach rate, collected during experimental sessions are used to analyze individual and group task performance as it was done for our initial publication on the RECORD system (Ibáñez Alcalá *et al*., 2024). In this paper, we analyze a new feature, ‘Time in reward zone’, which quantifies how long rat spends within the active reward zone during a trial.

### Psychometric function shape analysis

To analyze the psychometric function for each session of individual animals, we fitted the data using the function *f*(*x*) = *d* + (*a* − *d*)/(1 + (*x*/*c*)^*b*^). This function characterizes the sigmoidal relationship between the stimulus intensity, represented by the concentration of sucrose on the *X*-axis, and approach rate. In this equation, *a* and *d* represent the upper and lower horizontal asymptotes are represented, respectively, indicating the maximum and minimum response levels. *b* and *c* are the growth rate parameter and inflection point, respectively. For robust analysis, we only included sessions where individual animals completed at least 40 trials, disregarding any sessions with fewer trials. This threshold ensures the reliability of the fitted psychometric functions by providing sufficient data points for accurate curve fitting.

### Statistical analysis

All statistical analyses were performed using MATLAB (R2022a) software. The kernel probability density of inflection points between groups was compared using the two-sample Kolmogorov-Smirnov (KS) test. To address the issue of unbalanced sample sizes across different conditions (alcohol task performers vs. non-alcohol task performers) a non-parametric bootstrapping method with 1,000 iterations was employed during the KS test. Approach rates between groups were compared using multivariate analysis of variance (MANOVA). The F-test was conducted to determine whether the variances of two populations were equal. F-test results at four different reward levels were combined using Fisher’s method to obtain a combined p-value. A psychometric function for a session was classified as sigmoid if the coefficient of determination (*R*^2^) was ≥ 0.9. The ’fraction of sigmoid’ for an animal within a particular health group was calculated by dividing the number of sigmoid sessions by the total number of sessions. Most rats met or exceeded a threshold of 0.7, meaning more than 70% of sessions were sigmoidal. Rats who had under 70% of sessions that formed sigmoidal functions were considered to perform the task abnormally. A chi-square test was performed to assess whether there was a significant difference in the proportion of rats who were above versus below 0.7 between the two genders within the same health group.

## Results

### Alcohol alters psychometric functions

The psychometric functions of approach rate typically form a sigmoidal function (individual example **Fig. 1d**). During NCCB (**Fig. 1e**) and CCB (**Fig. 1f, top**), tasks performed by the rats before being introduced to alcohol, the individual psychometric functions generated during each are sigmoidal with the highest approach rates generally observed at 9% sucrose. During AA task performance, which lasted around three months and offered sucrose concentrations mixed with ethanol in inverse proportions (see materials and methods: **Acute alcohol task**), individual psychometric functions began to become less sigmoidal and seem to indicate increased approach rates for some individuals at 2% and 5% (**Fig. 1f**, bottom). We then compared the distribution of inflection points comparing NCCB, CCB, Non-conflict post alcohol (NCPA), and conflict post alcohol (CPA). The post-alcohol tasks were the same as NCCB and CCB tasks but performed for two months after AA task performance. The population distributions of the average inflection point (where the psychometric functions switch between approaching or avoiding 50% of the time) had no significant differences when comparing AA to NCCB or NCPA (**Fig. 1g,h**, both panels represent the same data, top) whereas the distribution is significantly left shifted comparing AA to CCB with no significant differences compared to CPA (**Fig. 1g,h**, both panels represent the same data, bottom, p < 0.0001 CCB to AA).

We then compared the average approach rates for each reward level during AA to NCCB (**Fig. 1i top**, MANOVA, p = 0.001), NCPA (**Fig. 1i top**, MANOVA, p = 0.05) and CCB (**Fig. 1i bottom**, MANOVA, p < 0.0001) increased approach rates during AA task performance. When averaging approach rates for each task we continue to see significantly higher approach rates compared to NCCB (**Supplementary Fig. 1a left,** MANOVA, NCCB compared to AA p = 0.001) and CCB (**Supplementary Fig. 1a bottom**, MANOVA, CCB compared to AA p < 0.0001). To identify whether heterogeneous populations exist, variance was plotted and found to be significantly different across AA, NCCB (**Fig 1j top**, F-test, p = 0.0096), CCB (**Fig 1j bottom**, F test, p < 0.0001), and CPA (**Fig 1j bottom**, F test, p < 0.0001).

### Sex differences to alcohol

Looking at individual functions formed during NCCB then AA split by sex it seems that males generate more abnormal psychometric functions during AA task performance than females (**Figs. 2a, b, top male, bottom female for all figures**). Examining the distribution of inflection points across males and females we see a significant left shift in the male distribution compared to NCCB (**Fig. 2c, top**, p = 0.004) while females had no significant changes in the inflection point distribution (**Fig. 2c, bottom**). This was corroborated by analyzing the cumulative density functions (**Fig. 2d**). Approach rates for males during AA task performance were significantly increased when compared to NCCB (**Fig. 2e, top**, **Supplemental Fig. 2c left**, p < 0.0001), while females had no significant differences across NCCB, NCPA, and AA (**Fig. 2e, bottom, Supplemental Fig. 2c, right**). Individual psychometric functions are similarly impacted across sex during AA task performance compared to CCB task performance (**Fig. 2f**). Similarly to NCCB and NCPA, the AA distribution was only significantly shifted in males across both density distributions and cumulative density functions (**Figs. 2g, h**, males AA compared to CCB, p < 0.0001). There was also significant variation in males between CCB and AA task performance, but no such significant variation was observed in females **(Supplemental Fig. 2f**, CCB vs AA: p < 0.0001 males, CCB vs. CPA: p < 0.0001 males).

### Alcohol alters time spent in reward zones

Evaluating alternative behaviors during task performance, we measured the time each rat spent in individual reward zones (**Supplemental Fig, 2a**). On average, rats performing AA spent more time in rewards zones containing 0.5%, 2% and 5% sucrose, but not 9%, compared to NCCB (**Supplemental Fig 2b top**, MANOVA, p = 0.0002) and CCB with a significant difference between CCB and CPA (**Supplemental Fig 2b bottom**, MANOVA, CCB compared to AA p < 0.0001, CCB compared to CPA p < 0.0001). Additionally, probability density plots of time spent in reward zones revealed that alcohol induced shifts significantly different from NCCB (**Supplemental Fig 2c top**, KS test, p = 0.003) and CCB (**Supplemental Fig 2c bottom**, KS test, p = 0.004). When split across genders, males spend significantly more time in reward zones (**Supplemental Fig 2e top**, MANOVA, p < 0.0001) and significantly altered probability density shifts (**Supplemental Fig 2f top**, KS test, p = 0.005) when completing an AA versus NCCB. Similar trends are observed when males, but not females, demonstrate increased time in reward zones during CCB task performance (**Supplemental Fig 2h top**, MANOVA, p < 0.0001) when performing AA compared to CCB. Males also (**Supplemental Fig 2i top**, KS test, p = 0.009) while there were no significant differences between distributions in females (**Supplemental Fig. 2i**, bottom).

### Identifying abnormal task performance in individuals using psychometric functions

The psychometric functions formed during behavioral task performance across sessions may be a helpful marker for identifying individuals who are impacted by alcohol presentation and consumption (**Supplemental Fig. 3a**). Across all tasks, animals are considered “abnormal” when the proportion of sigmoidal functions formed is less than 70% (which is the same standard we used for our initial publication where we develop and validate the RECORD system(Ibáñez Alcalá *et al*., 2024)) of non-alcohol conditions, determined using the chi-square (**Fig. 3a**). Non-abnormal animals, in contrast, seem to maintain more consistent sigmoid-shaped functions across task types (**Fig. 3b**). We identified 7 abnormal males (**Fig. 3c left**) and 2 abnormal females (**Fig. 3c right**). This difference between the sexes was determined to be significant using a chi-square test (**Fig. 3d,** p=0.025). Further, only one male and 3 females had less than 70% of sigmoidal sessions during both NCCB (**Supplemental Fig. 3b-c**) and CCB tasks (**Supplemental Fig. 3d-e**).

### Alcohol effects the proximal cost benefit tasks

Comparing psychometric functions of proximity tasks (**Fig 4a**) to tasks completed soon after alcohol task performance (**Fig 4b**), stark individual differences can be observed. Evaluating probability density plots across each stage of non-conflict cost benefit task, no significant difference is observed. When comparing probability density plots between each stage of conflict cost benefit task, a significant difference in peaks between CCB and PC (**Fig 4c bottom**, KS test, p < 0.0001) is observed. When comparing NCCB to PNC (MANOVA, p= 0.0023) and NCPA (MANOVA, p=0.046), significantly different approach rates (**Fig. 4e top**) were observed. While approach rates for each stage of this task follow an overall trend of increased acceptance as reward increases, only NCCB compared to PNC retain significant variability (**Fig 4f top**, F-test, p=0.0015). Evaluating approach rate across each stage of the conflict cost benefit task, both PC (**Fig 4d bottom**, MANOVA, p<0.0001) and CPA (**Fig 4d bottom**, MANOVA, p=0.0413) were significantly different from CCB. Variance was also significantly different when comparing PC (**Fig 4e bottom**, F-test, p<0.0001) and CPA (**Fig 4e bottom**, F-test, p<0.0001) to CCB.

### Sex differences are observed during the proximal tasks

Comparison of individual psychometric functions, separated by sex, across PNC (**Fig 5a**) and NCPA (**Fig 5b**). There were no significant differences in inflection point distributions for PNC task performance compared to NCCB and NCPA (**Fig 5c**). Males also demonstrate shifted approach rates in PNC (**Fig 5d top**, MANOVA, p<0.0001), while females do not. Comparisons between NCCB and PNC display significantly different variance in both male (**Fig 5e top**, F-test, p=0.015) and females (**Fig 5e bottom**, F-test, p=0.0153). Individual psychometric functions from PC (**Fig 6f left**) and CPA (**Fig 5g right**) can be evaluated based on sex. Strikingly, both males (**Fig 5h top**, KS test, p=0.002) and females (**Fig 5h bottom**, KS test, p=0.02) demonstrate significant alterations in density when comparing CCB and PC tasks. Notably, males (**Fig 5j top**, MANOVA, p<0.0001) and females (**Fig 5j bottom**, MANOVA, p=0.005) exhibit alterations in approach rate during proximal cost benefit conflict task.

### Time spent in reward zone altered in the proximal tasks

Proximal non-conflict task demonstrates significantly increased time spent in reward zones for 0.5, 2, and 5% sucrose (**Supplemental Fig 4a** top, MANOVA, p=0.0209). However, both PNC (**Supplementary Fig 4b top**, KS test, p=0.0000) and NCPA (**Supplementary Fig 4b top**, KS test, p=0.0350) tasks display significant alterations in density when compared to NCCB. Proximal (**Supplementary Fig 4a bottom**, MANOVA, p=0.0001) and post alcohol (**Supplemental Fig 4a bottom**, MANOVA, p=0.0001) cost benefit conflict tasks demonstrate significantly increased time spent in reward zones containing 0.5%, 2%, and 5% sucrose compared to CCB. When evaluating gendered effects of time spent in reward zones, males but not females demonstrated increased time spent in 0.5%, 2%, and 5% sucrose during both PNC (**Supplementary Fig 4c top**, MANOVA, p=0.0021) and NCPA (**Supplementary Fig 4c top**, MANOVA, p=0.047). Males also exhibit increased density in PNC (**Supplemental Fig 4d top**, KS test, p=0.003) but not NPA tasks. This trend of increase time spent in 0.5%, 2%, and 5% sucrose conserved, and male dominated in PC (**Supplemental Fig 4e top**, MANOVA, p<0.0001) and CPA (**Supplemental Fig 3e bottom**, MANOVA, p<0.0001) tasks. This is further supported by the significantly altered density peaks in males performing PC (**Supplemental Fig 4f top**, KS test, p<0.0001) and CPA (**Supplemental Fig 4f top**, KS test, p<0.0001).

### Identification of continued non-sigmoidal function during the proximal tasks

Interested in determining whether abnormal populations in proximal non-conflict and conflict tasks were gender exclusive, a chi squared test was used to quantify a 70% difference in fraction of a sigmoid. Notably, half the males in PNC displayed behavioral abnormalities while all females displayed did not (**Fig 6a-b**, chi-square test, p=0.0098). When comparing abnormal males and females in the context of NCPA, neither group was significant (**Supplementary Fig 4g-h**). During PC, males displayed greater instances of abnormalities (n=5) than females (n=2), but the two were not statistically different (**Fig 6c-d**, chi-square test, p=0.16). Interestingly, in CPA both males and females exhibit equal numbers of abnormal individuals (**Supplemental Fig 4i-j**, chi-square test, p= 1.0).

## Discussion

Overall, we introduced alcohol in inverse concentrations to an approach-avoid trade-off task. By presenting sucrose and alcohol in inverse concentrations (highest sucrose concentration with lowest alcohol concentration), rodents could minimize their exposure to alcohol while still attaining the attractive sucrose reward. However, we saw that rats had significantly different approach rates across various features once alcohol became integrated into the task environment (**Fig. 1**). When looking at approach rates for each level of reward the change in approach rate is driven by a higher approach rate for 2% and 5% sucrose, the moderate sucrose and alcohol combinations (**Supplemental Fig. 1**) as opposed to high alcohol (0.5%) or high sucrose concentration (9%).

When we looked at sex differences after introducing alcohol into the task environment, we found that males were significantly affected by the presence of alcohol, shifting their approach rates towards 2% and 5% while females were minimally affected (**Fig. 2**). One potential reason males start approaching 2% and 5% more while decreasing their approach for 9% could be that they view moderate sucrose with moderate alcohol more rewarding than either on their own, like how some people prefer mixed drinks or cocktails as opposed to strong spirits or soda.

Alcohol also significantly increased the time spent in reward zones; this significance is also mostly driven by males (**Fig. 3**). We then used behavioral psychometric functions to try and delineate between rats that presented abnormal behaviors during the alcohol task and rats that did not. We looked at the proportion of sessions that were sigmoidal vs those that were not and identified seven males and three females that had less than 70% of sessions that were sigmoidal (**Fig. 4**). Looking at task performance after alcohol consumption we saw that when compared to CCB and NCCB before acute alcohol, PC and PNC are significantly altered with the preference for 2% and 5% being retained while the preference for 9% occasionally dips below pre-alcohol levels (**Fig. 5,6**). We also found that markers of acute alcohol abnormality persist for five male rats and two female rats after alcohol task performance (**Fig. 7**).

In the United States, half of adults reported consuming alcohol in the thirty days prior to being surveyed, 17% of adults reported binge drinking, and 6% reported drinking heavily (Centers for Disease Control and Prevention (CDC), 2023). Alcohol abuse is characterized by persistent, habitual drinking despite social, medical, and economic decline. Interestingly, despite how prevalent drinking alcohol is, most individuals who drink do not develop an alcohol use disorder (Kranzler, 2023). Researchers, however, often face the difficulty of developing an ethologically relevant rodent alcohol task since alcohol is aversive to rodents (Pautassi *et al*., 2011).

Many other rodent paradigms looking at alcohol use and decision-making examine binge drinking patterns or chronic alcohol consumption(Strong *et al*., 2010; Jury *et al*., 2017). Additionally, these conditions are typically done using a water and ethanol mixture without sucrose(Crabbe *et al*., 2009; Hwa *et al*., 2011). These studies generally find that female rodents are the ones that consume more sucrose(Strong *et al*., 2010; Hwa *et al*., 2011; Jury *et al*., 2017). In our task, we found that presenting a sucrose-alcohol mixture into the maze caused a significant change in how males perform the task, where they approached 2% and 5% sucrose more than in the non-alcohol task conditions. Females in our task did not demonstrate many significant differences whether alcohol was present or not. Overall, this research, potentially due to the approach-avoid style task, mixture of alcohol with sucrose, and acute nature of the alcohol exposure finds a greater shift in male decision-making and behavior with minimal shifts observed in female rodents.

This aversion has required the use of various approaches like forcing alcohol consumption (Thiele and Navarro, 2014; Mendoza-Ruiz *et al*., 2018) or breeding animals genetically predisposed to alcohol consumption (Timme *et al*., 2020; Borruto *et al*., 2021; Sauton *et al*., 2021). Traditional forced alcohol models are effective in representing alcohol use disorder but are less suited for mimicking acute/social drinking. We sought to mimic acute drinking by offering differing concentrations of alcohol and using a paradigm where the alcohol can be approached (accepted) or avoided (rejected). We utilized our RECORD task environment to design a battery of tasks that incorporate elements commonly used including sweetened alcohol, cost-benefit trade-offs, and run-away paradigms. A notable strength of the RECORD system is the ability to offer a “gradient” of trade-offs within a single session.

Future work exploring acute alcohol consumption could use a similar task setup with *in vivo* recordings to uncover correlated neural activity as the task is performed. Exploring these neuronal recordings and other alcohol factors, like delivering chronic alcohol, could give further insight into the physiological activities underlying acute/social alcohol consumption and how it may differ in individuals with alcohol use disorder. Additionally, further exploration into identifying individuals who are more impaired by alcohol through the alterations in psychometric functions and associated neural correlates could help identify at risk groups which may eventually lead to earlier interventions for those predisposed to alcohol use disorder.

## Data Availability

The data used within this manuscript is available at https://dataverse.harvard.edu/dataset.xhtml?persistentId=doi:10.7910/DVN/YVPYUJ&version=DRAFT

## Code Availability

All codes used for this manuscript can be accessed with the following link: https://github.com/atanugiri/Alcohol-Project/tree/main

## Ethics declarations

The authors declare no competing interests.

## Animal ethics statement

The project received approval for all protocols from the University of Texas at El Paso Institutional Animal Care and Use Committee and followed the Guide for Care and Use of Laboratory Animals (IACUC reference number: A-202009-1).

## Acknowledgements

We appreciate Dr. Aguilera who funded several students who worked on this project through U-RISE T34GM145 and G-RISE T32GM144919.

---

**Supp Fig 1:**
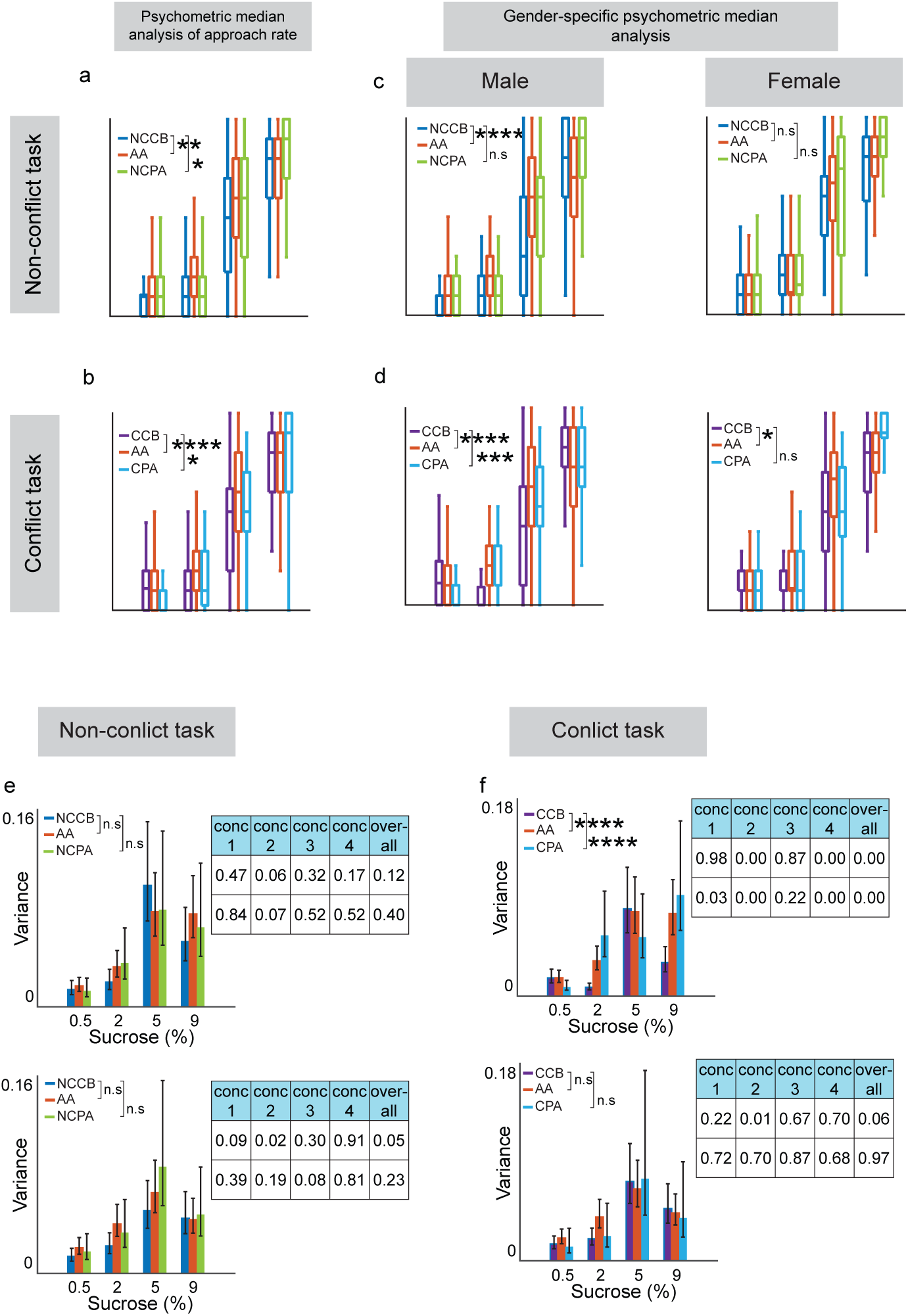
Psychometric functions altered following alcohol. (**a**) Difference in average approach rates is significant between NCCB and AA tasks (NCCB vs AA MANOVA, p = 0.001, NCCB vs NCPA, p = 0.05). (**b**) Differences between CCB vs. CPA and AA are significant (CCB vs AA, MANOVA, p < 0.0001, CCB vs CPA, p = 0.04). (**c**) sex differences in approach during non-conflict tasks were only present in males NCCB vs AA (MANOVA, p < 0.0001). (**d**) During conflict tasks, male approach rate was significantly different between CCB vs AA (p < 0.0001) and CPA (p = 0.0007) while females had significantly different approach rates between CCB and AA (p = 0.01). (**e**) Variance in approach rate during non-conflict task is insignificant across both sexes. (**f**) Variance is significant during conflict tasks in males (p < 0.0001 for both CCB vs AA and CCB vs CPA) but not females.

**Supplemental Fig 2:**
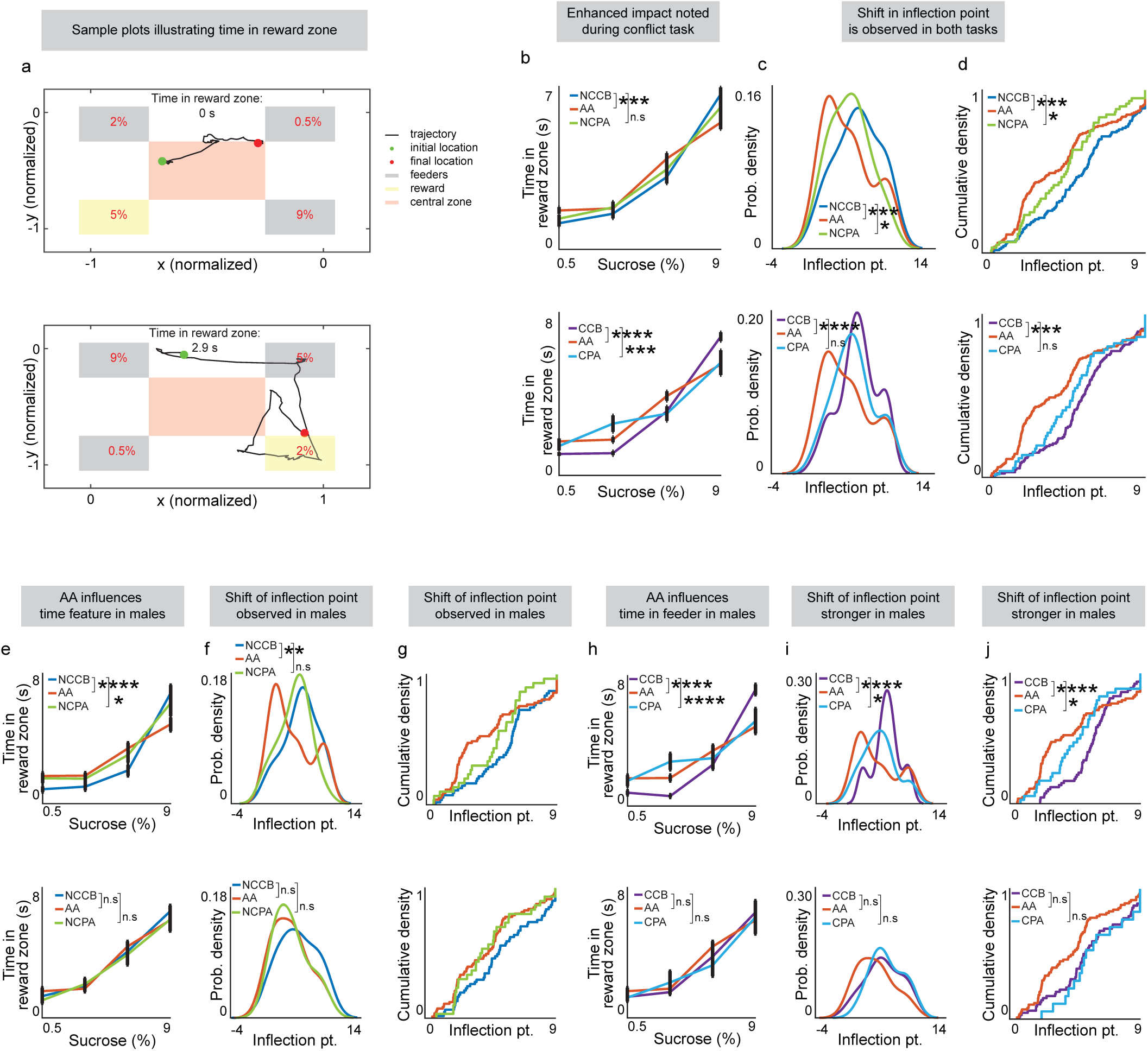
Alcohol alters time spent in reward zones. (**a**) Example plots of individual trials demonstrating time in reward zone across four sucrose concentrations. (**b**) Significantly different average time in reward zones between NCCB and AA tasks (MANOVA, p = 0.0002, NCCB vs NCPA, p = 0.05), and CCB and AA tasks (MANOVA, p < 0.0001). (**c**) Inflection point is significantly different between NCCB and AA (KS test, p = 0.003) and CCB and AA tasks (KS test, p = 0.004). (**d**) Inflection point distribution represented as a cumulative density function. (**e**) Time in reward zone is significantly different between NCCB and AA tasks for males only (MANOVA, p < 0.0001). (**f**) Inflection point is significantly different in male NCCB and AA tasks (KS test, p = 0.005). (**g**) Inflection point distribution represented as a cumulative density function. (**h**) Males display significantly different times spent in reward zones in CCB and AA (MANOVA, p < 0.0001). (**i**) Males (KS test, p = 0.009) are also marked by significantly altered inflection points in CCB compared to AA. (**j**) Inflection point distribution represented as a cumulative density function.

**Supp Fig 3:**
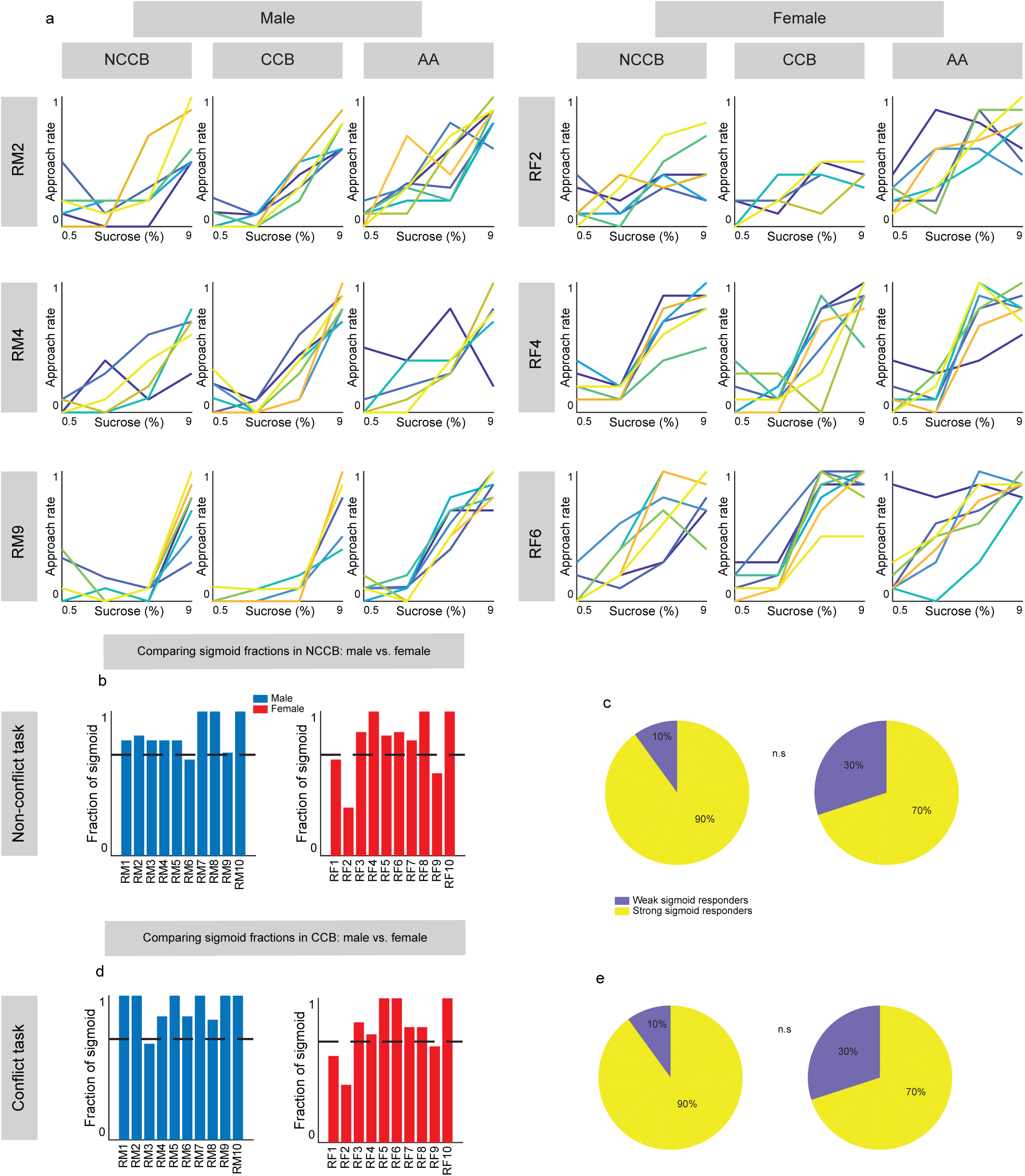
Normal versus abnormal task performance in individuals following alcohol. (**a**) Individual male and female psychometric profiles in NCCB, CCB, and AA tasks. (**b**) Two plots showing the fraction of sigmoid across all sessions in NCCB for individual males (n = 10) and females (n = 10), where there is no significant difference in (**c**) abnormality across genders (chi-square test, p = 0.2636). (**d**) Two plots showing the fraction of sigmoid across all sessions in CCB for individual males (n = 10) and females (n = 10), where there is no significant difference in (**e**) abnormality across genders (chi-square test, p = 0.26).

**Supp Fig 4:**
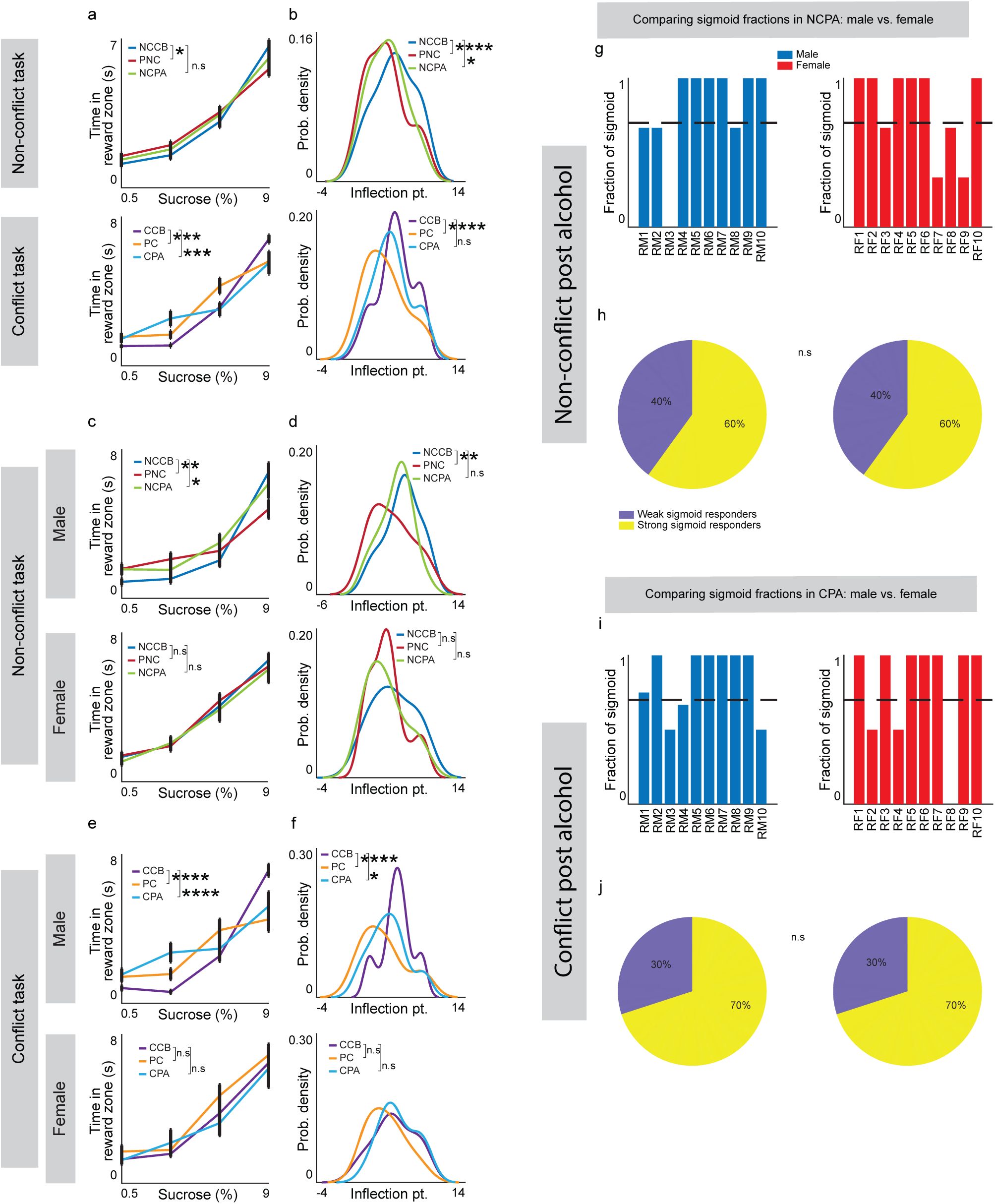
Alcohol affects time and individual differences in non-alcohol related decision-making. (**a**) Significantly different in time in reward zone in NCCB vs PNC (MANOVA, p = 0.0209), but not in NCCB vs NCPA (MANOVA, p = 0.6699). Both PC (MANOVA, p = 0.0001) and CPA (MANOVA, p = 0. 0001) tasks are significantly different from the CCB task. (**b**) Changes in inflection point are more prominent in NCCB vs PNC (KS test, p = 0.0000) than in NCCB vs NCPA (KS test, p = 0.0350). (**c**) Only CCB vs PC show a significant alteration in inflection point (KS test, p = 0.0000). PNC (MANOVA, p = 0.0021) and NCPA (MANOVA, p = 0.0474) compared to NCCB show a significant difference in time spent in reward zone across males only. (**d**) Only male PNC vs NCCB show a statistical difference in inflection point of kernel probability densities (KS test, p = 0.0030). (**e**) Males show significant difference in time spent in reward zones for PC (MANOVA, p = 0.0000) and CPA (MANOVA, p = 0.000) tasks, while females are insignificant. (**f**) Probability densities are significantly different in male PC (KS test, p = 0.0000) and CPA (KS test, p = 0.0310) tasks, but females do not demonstrate significance. (**g**) Two plots showing the fraction of sigmoid across all sessions of NCPA for individual males (n = 10) and females (n = 10), where neither gender has a significant number (chi-square test, p = 1.0000) of (**h**) abnormal animals. (**i**) Two plots showing the fraction of sigmoid across all sessions of the CPA task for individual males (n = 10) and females (n = 10), (**j**) where the number of animals having sigmoidal and non-sigmoidal psychometric profiles for both genders are not significant (chi-square test, p = 1.0000).

